# Identification of HLA-A*24:02-restricted CTL candidate epitopes derived from the non-structural polyprotein 1a of SARS-CoV-2 and analysis of their conservation using the mutation database of SARS-CoV-2 variants

**DOI:** 10.1101/2021.09.21.461322

**Authors:** Akira Takagi, Masanori Matsui

## Abstract

COVID-19 vaccines are currently being administrated worldwide and playing a critical role in controlling the pandemic. They have been designed to elicit neutralizing antibodies against Spike protein of the original SARS-CoV-2, and hence they are less effective against SARS-CoV-2 variants with mutated Spike than the original virus. It is possible that novel variants with abilities of enhanced transmissibility and/or immunoevasion will appear in the near future and perfectly escape from vaccine-elicited immunity. Therefore, the current vaccines may need to be improved to compensate for the viral evolution. For this purpose, it may be beneficial to take advantage of CD8^+^ cytotoxic T lymphocytes (CTLs). Several lines of evidence suggest the contribution of CTLs on the viral control in COVID-19, and CTLs target a wide range of proteins involving comparatively conserved non-structural proteins. Here, we identified twenty-two HLA-A*24:02-restricted CTL candidate epitopes derived from the non-structural polyprotein 1a (pp1a) of SARS-CoV-2 using computational algorithms, HLA-A*24:02 transgenic mice and the peptide-encapsulated liposomes. We focused on pp1a and HLA-A*24:02 because pp1a is relatively conserved and HLA-A*24:02 is predominant in East Asians such as Japanese. The conservation analysis revealed that the amino acid sequences of 7 out of the 22 epitopes were hardly affected by a number of mutations in the Sequence Read Archive database of SARS-CoV-2 variants. The information of such conserved epitopes might be useful for designing the next-generation COVID-19 vaccine that is universally effective against any SARS-CoV-2 variants by the induction of both anti-Spike neutralizing antibodies and CTLs specific for conserved epitopes.

**Importance:** COVID-19 vaccines have been designed to elicit neutralizing antibodies against the Spike protein of the original SARS-CoV-2, and hence they are less effective against variants. It is possible that novel variants will appear and escape from vaccine-elicited immunity. Therefore, the current vaccines may need to be improved to compensate for the viral evolution. For this purpose, it may be beneficial to take advantage of CD8^+^ cytotoxic T lymphocytes (CTLs). Here, we identified twenty-two HLA-A*24:02-restricted CTL candidate epitopes derived from the non-structural polyprotein 1a (pp1a) of SARS-CoV-2. We focused on pp1a and HLA-A*24:02 because pp1a is conserved and HLA-A*24:02 is predominant in East Asians. The conservation analysis revealed that the amino acid sequences of 7 out of the 22 epitopes were hardly affected by mutations in the database of SARS-CoV-2 variants. The information might be useful for designing the next-generation COVID-19 vaccine that is universally effective against any variants.

## Introduction

The severe acute respiratory syndrome coronavirus 2 (SARS-CoV-2) is the causative agent of the coronavirus disease 2019 (COVID-19) pandemic, which has resulted in more than 222 million infections and 4.6 million deaths around the world as of 10th September 2021. To bring the pandemic under control, a number of vaccine candidates are being developed at an unprecedented speed, and several of them are currently being administrated all over the world. These include two mRNA vaccines of BNT162b2 (Pfizer/BioNTech) and mRNA-1273 (Moderna), and adenoviral-vectored vaccines such as ChAdOx1 nCoV-19 (AstraZeneca), Gam-COVID-Vac (Sputnik V, Gamaleya Research Institute), and Ad26.COV2.S (Janssen). Most of these vaccines have been designed to elicit neutralizing antibodies against the SARS-CoV-2 spike (S) protein that block the interaction between SARS-CoV-2 and the angiotensin-converting enzyme 2 on target cells, and thereby preventing SARS-CoV-2 infection. The Pfizer/BioNTech and Moderna mRNA vaccines showed 95% and 94.1% efficacy in preventing the onset of disease caused by SARS-CoV-2 (1, 2), respectively, whereas adenoviral-vectored vaccines demonstrated protection at a slightly lower but sufficient efficacy (3, 4). Therefore, it was initially speculated that these vaccines would put an end to the pandemic sooner or later. However, the recent emergence of various SARS-CoV-2 variants has made us realize that the initial idea was optimistic.

Although SARS-CoV-2 changes more slowly than most other RNA viruses because of a proofreading mechanism (5), its variants have continuously emerged in the circulating viruses. Since the fall of 2020, variant strains with enhanced transmissibility have been found in the United Kingdom (Alpha or B.1.1.7), South Africa (Beta or B.1.351) and Brazil (Gamma, B.1.1.28 or P.1). Because the vaccines were directed against the original SARS-CoV-2 virus that appeared in 2019, it was questioned whether the vaccines would quell SARS-CoV-2 variants. Although the impact of the first detected Alpha variant on anti-SARS-CoV-2 humoral immunity was found to be moderate (6–8), the Beta and Gamma variants significantly reduced susceptibility to neutralizing antibodies (8, 9). Particularly, the Beta was the most resistant to available monoclonal antibodies, convalescent and vaccinated sera (7, 8, 10–13). Fortunately, however, it was demonstrated that the BNT162b2 (Pfizer/BioNTech) and mRNA-1273 (Moderna) mRNA vaccines were highly effective against the Beta variant infection in Qatar, and people who had received two doses of this vaccine were almost completely protected from severe disease caused by the Beta variant (14, 15), suggesting that the current vaccines are still effective even against the Beta variant. In December 2020, another variant of concern, the Delta strain (B.1.617.2) has appeared in India. The Delta variant is more transmissible than the highly contagious Alpha variant (16), and this fastest strain has been expected to rapidly outcompete other variants and become the dominant lineage in many parts of the world (17). Furthermore, it was reported that viral loads in Delta infections were ∼1,000 times higher than those in initial infections in early 2020 (18). It was also shown that unvaccinated individuals infected with the Delta were more likely to be hospitalized than unvaccinated people infected with the Alpha (19) although it is still unclear whether the Delta variant causes more severe illness than the previous strains. On the other hand, real-world data demonstrated that only modest differences in the vaccine efficiency were observed between the Delta and the Alpha after the receipt of two doses of vaccine (BNT162b2 or ChAdOx1 nCoV-19) (20), suggesting that two doses of the current vaccine may be effective against the Delta. However, the breakthrough infection with the Delta is often observed in fully vaccinated individuals. It has been proven that the vaccinated people with the Delta breakthrough infection may not develop severe disease but may have the potential to transmit SARS-CoV-2 to others as the same rates as those who are unvaccinated (21). Taken together, the current vaccines are likely to be still efficient for existing variants including the Beta and Delta, but becoming less effective against the variants than the original virus. In addition to these four variants of concern, several variants of interest have emerged all over the world. Accordingly, it is possible that new variants with abilities of more enhanced transmissibility and/or immunoevasion will appear in the near future and perfectly escape from natural and vaccine-elicited immunity. Therefore, the current vaccines may need to be improved to the next-generation vaccines in order to compensate for the viral evolution.

In general, CD8^+^ cytotoxic T lymphocytes (CTLs) play a crucial role for the clearance of virus as well as neutralizing antibodies in the viral infection. CTLs recognize virus-derived peptides in association with major histocompatibility complex class I (MHC-I) molecules on the surface of antigen presenting cells and kill virus-infected target cells. In COVID-19, there was a greater proportion of SARS-CoV-2-specific CD8^+^ T cells in mild disease compared with severe case (22–25), suggesting a potential protective role of CD8^+^ T cell response. In fact, two persons with X-linked agammaglobulinemia recovered from pneumonia caused by the SARS-CoV-2 infection (26). In the virus-challenge experiment using rhesus macaques, depletion of CD8+ T cells in convalescent macaques that had been infected with SARS-CoV-2 partially abrogated the protective efficacy of natural immunity against rechallenge with SARS-CoV-2 (27), suggesting CD8^+^ T cells can contribute to virus control in COVID-19. The current mRNA vaccine and adenoviral-vectored vaccine elicit SARS-CoV-2 S protein-specific CD8^+^ CTLs as well as anti-S neutralizing antibodies (28), which might make these vaccines more efficient than inactivated and subunit vaccines. It is known that BNT162b2 mediates protection from severe disease as early as 10 days after prime vaccination, when neutralizing antibodies are hardly detectable. Since functional S-specific CD8^+^ T cells were shown to be already present at this early stage, CD8^+^ T cells were speculated to be the main mediators of the protection (29). Thus, several lines of evidence suggest the contribution of CTLs on the viral control in COVID-19, and therefore it may be beneficial to take advantage of CD8^+^ CTLs for the development of the next-generation vaccine. In addition, CTLs can target a wide range of proteins involving comparatively conserved non-structural proteins. A novel vaccine with ability to elicit conserved epitope-specific CTLs may not be affected by mutations of various SARS-CoV-2 variants.

As shown in Fig. 1, the 5’-terminal two-thirds of the genome of SARS-CoV-2 are composed of the open reading frame 1a (ORF1a) and ORF1b. The ORF1a encodes the polyprotein 1a (pp1a) which is a largest protein composed of 11 non-structural regulatory proteins (nsp1-11) in SARS-CoV-2. Due to its large size, it seems highly possible to find dominant epitopes in the pp1a. Saini et al. revealed that most of the immunodominant epitopes they identified belonged to the ORF1 region (30). In addition, it may be possible to identify conserved CTL epitopes in the pp1a because the ORF1 region is highly conserved within coronaviruses relative to structural proteins (31). From the above, we here attempted to identify conserved CTL epitopes derived from pp1a of SARS-CoV-2 using MHC-I transgenic mice. We focused on HLA-A*24:02-resctricted CTL epitopes because HLA-A*24:02 is relatively predominant in East Asians such as Japanese (32). This information might be useful for designing the next-generation COVID-19 vaccine that is universally effective against any SARS-CoV-2 variants by the induction of both anti-Spike neutralizing antibodies and CTLs specific for conserved epitopes.

**FIG 1.**
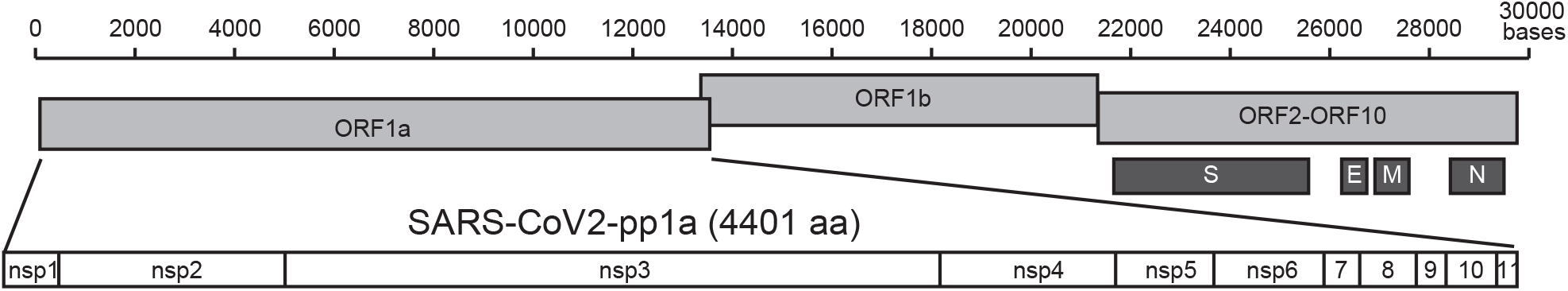
The linear diagrams of the SARS-CoV-2 genome and the protein subunits of ORF1a. The SARS-CoV-2 genome consists of ORF1a, ORF1b, and ORF2-ORF10. S, E, M, and N represent spike, envelope, membrane, and nucleocapsid, respectively. The ORF1a polyprotein (pp1a) is composed of eleven non-structural proteins, nsp1-nsp11.

## Results

### Prediction of HLA-A*24:02-restricted CTL epitopes derived from SARS-CoV-2 pp1a

To predict HLA-A*24:02-rescricted CTL epitopes derived from SARS-CoV-2 pp1a, we used a T-cell epitope database, SYFPEITHI (33). The top 80 epitopes in the database were selected and were synthesized into 9-mer peptides (Table 1). These epitopes were also evaluated by other three programs, IEDB (34), ProPred-1 (35), and NetCTL (36) (Table 1). Scores of the 80 epitopes in the four programs were assessed by classifying into four ranks (A: Excellent; B: Very good; C: Good; D: Poor) (SYFPEITHI: A ≥ 22, 20 ≤ B ≤ 21, 18 ≤ C ≤ 19, D = 17; IEDB: A < 0.1, 0.1 ≤ B < 0.5, 0.5 ≤ C < 1, D ≥ 1; ProPred-I: A ≥ 160, 50 ≤ B < 160, 20 ≤ C < 50, C < 20; NetCTL: A ≥ 1.70, 1.19 ≤ B < 1.70, 0.90 ≤ C < 1.19, C < 0.90) (Table 1). As shown in Table 1, the rank of each epitope was not always the same in the four programs, suggesting that multiple programs are needed to successfully predict CTL epitopes.

**TABLE 1.**
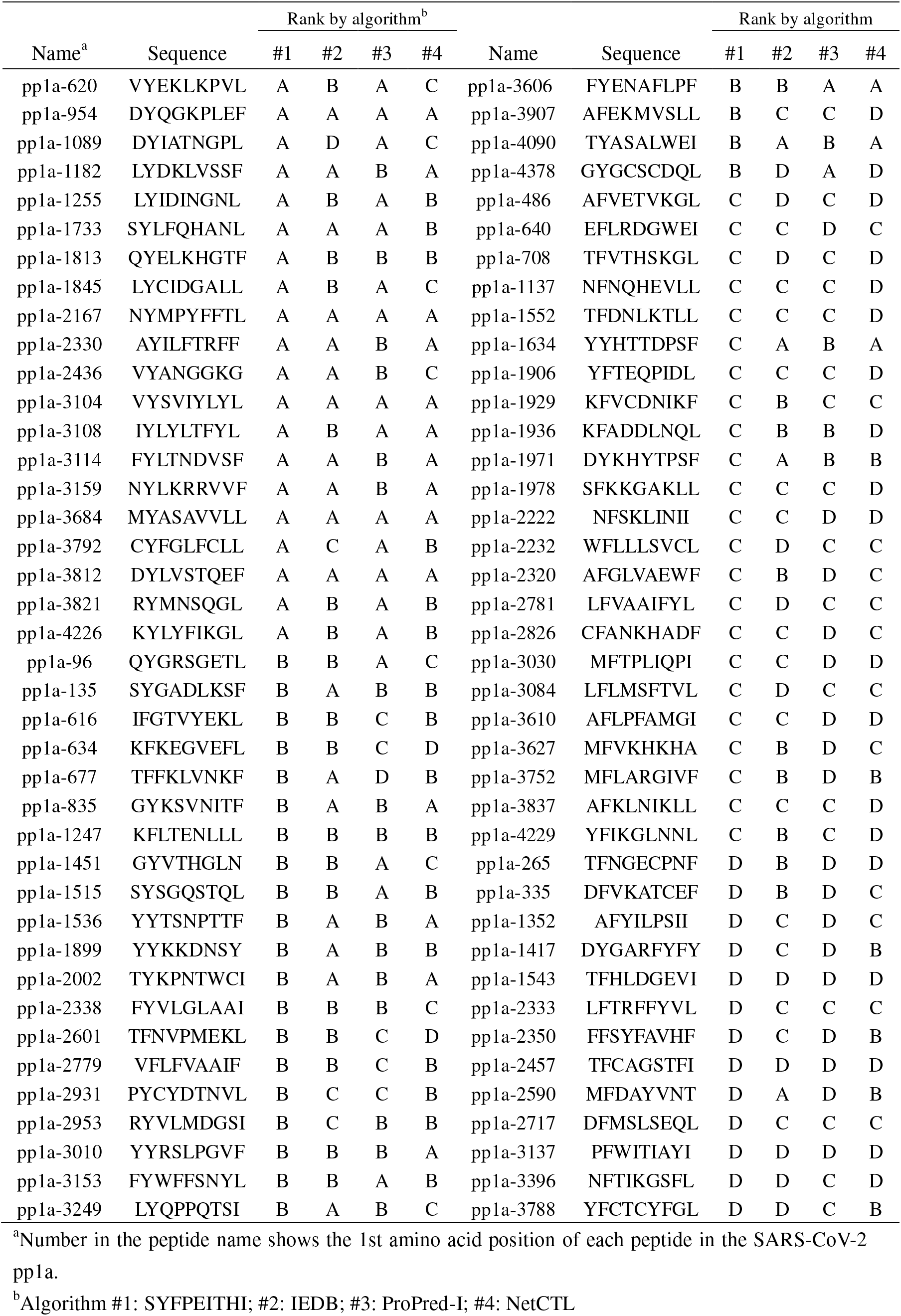
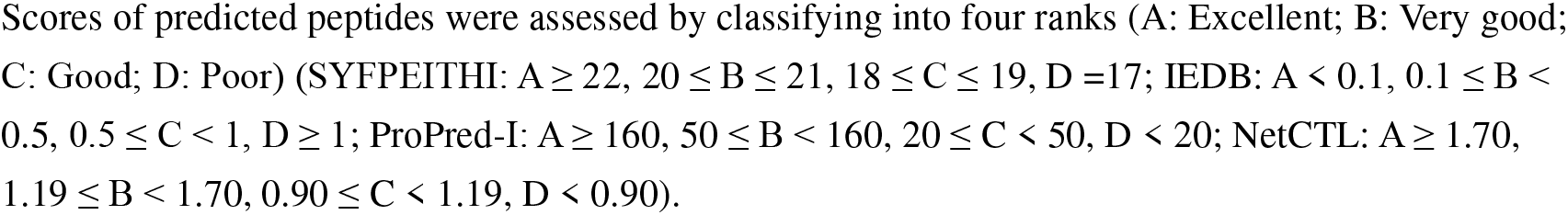
HLA-A*24:02-restricted CTL candidate epitopes for the SARS-CoV-2 pp1a

Eighty peptides were investigated for their binding affinities to HLA-A*24:02 molecules using TAP2-deficient RMA-S-HHD-A24 cells. Since the half-maximal binding level (BL_50_) value of a positive control peptide, Influenza PA_130-138_ (37) was 2.4 μM, we defined an extremely high binder with a BL_50_ value below 1.0 μM, a high binder with a BL_50_ value ranging from 1 to 10 μM, a medium binder with a BL_50_ value ranging from 10 to 80 μM, and a low binder with a BL_50_ value above 80 μM. Among 80 peptides, 11peptides and 10 peptides were extremely high binders and high binders, respectively, while 15 peptides were medium binders (Table 2). The remaining 44 peptides demonstrated low binding affinities or no binding to HLA-A*24:02 (Table 2). Comparison of the peptide binding affinity and the peptide rank in the 4 algorithms (Table 3) revealed that A-ranked peptides did not always show the high level of the peptide binding affinity to HLA-A*24:02. On the other hand, none of D-ranked peptides were classified into the extremely high group. When comparing the four programs in the prediction of extremely high binders and high binders, the IEDB program was likely to estimate them most accurately (Fig. 2).

**FIG 2.**
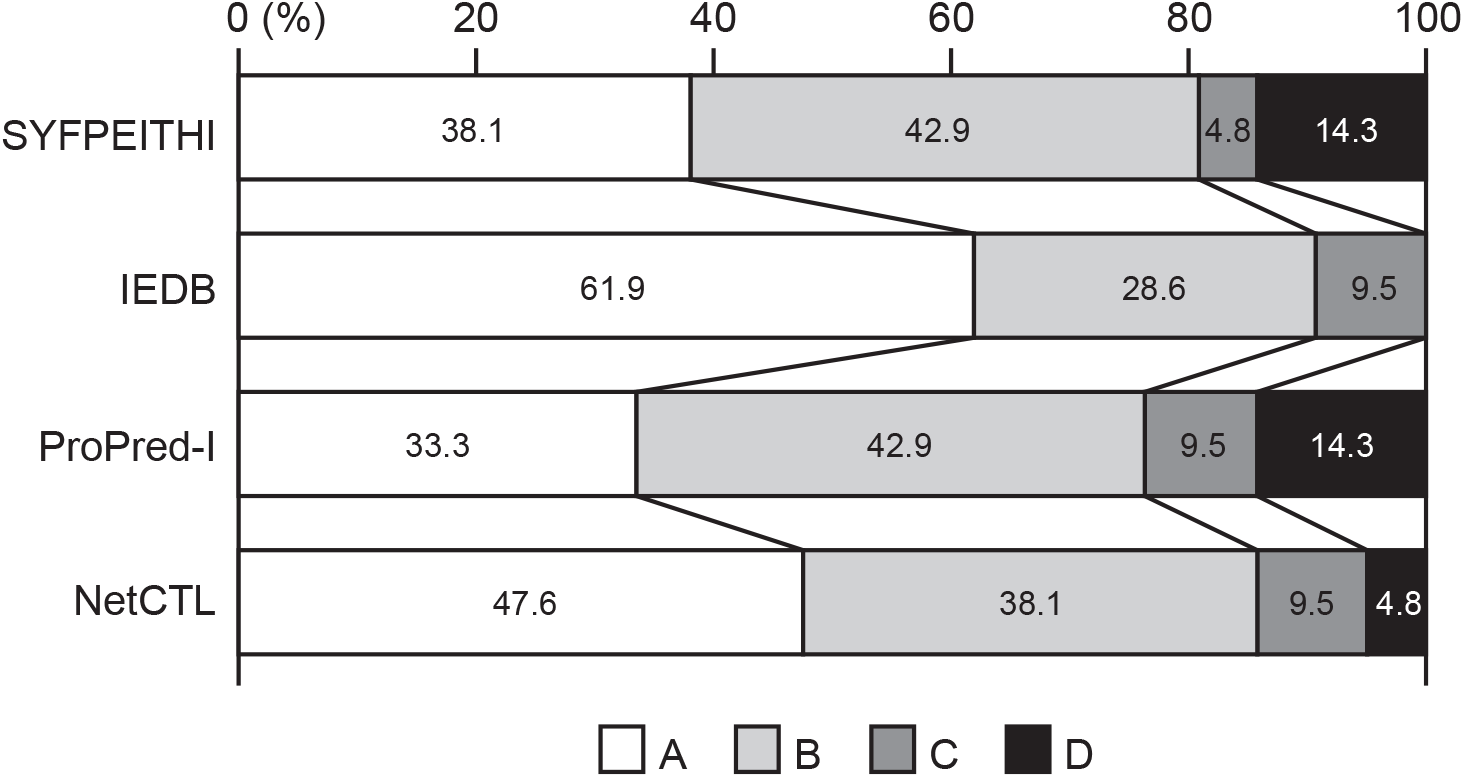
Percentages of peptides ranked from A to D (A: Excellent; B: Very good; C: Good; D: Poor) by each algorithm in the sum of extremely high peptides and high binder peptides.

**TABLE 2.**
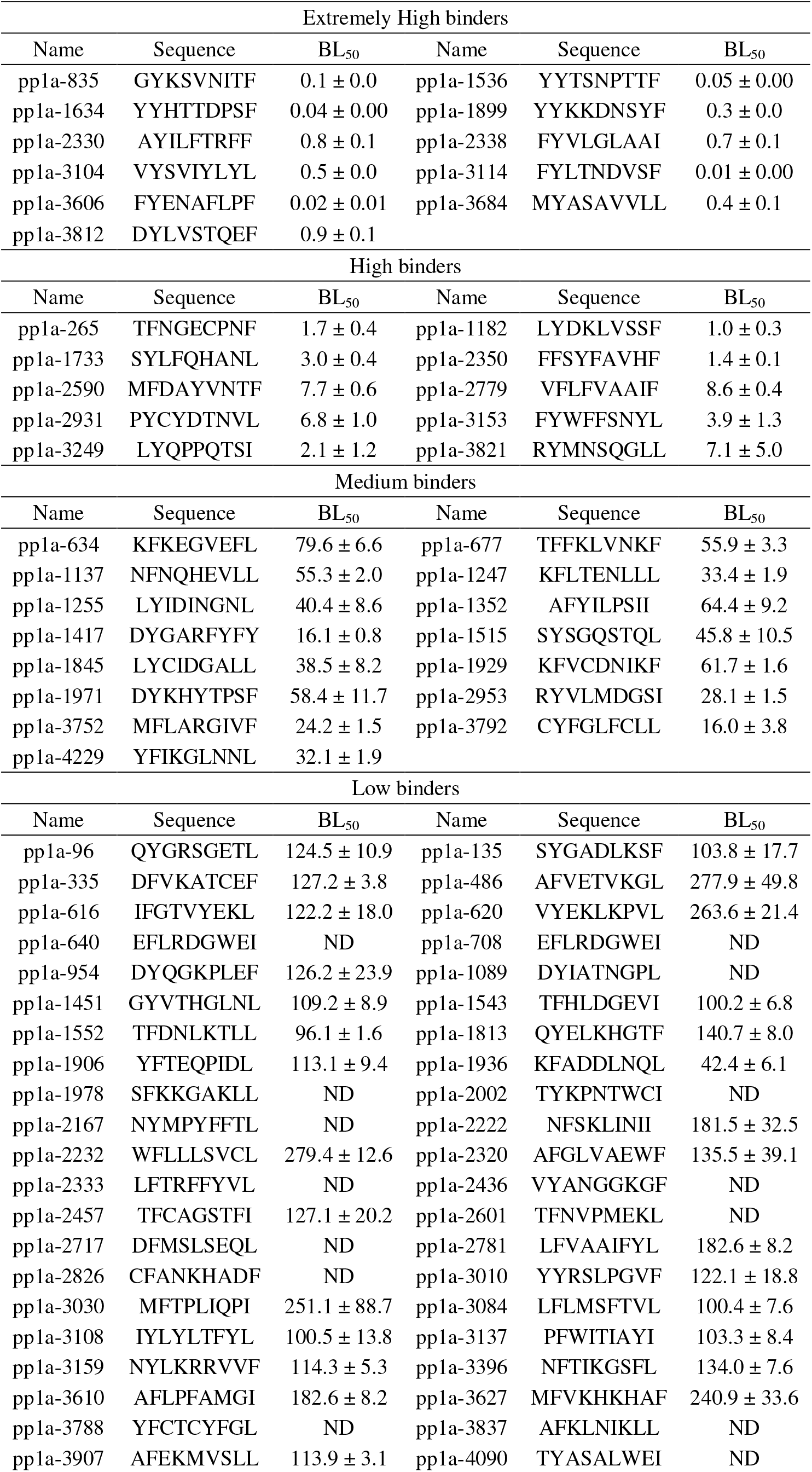
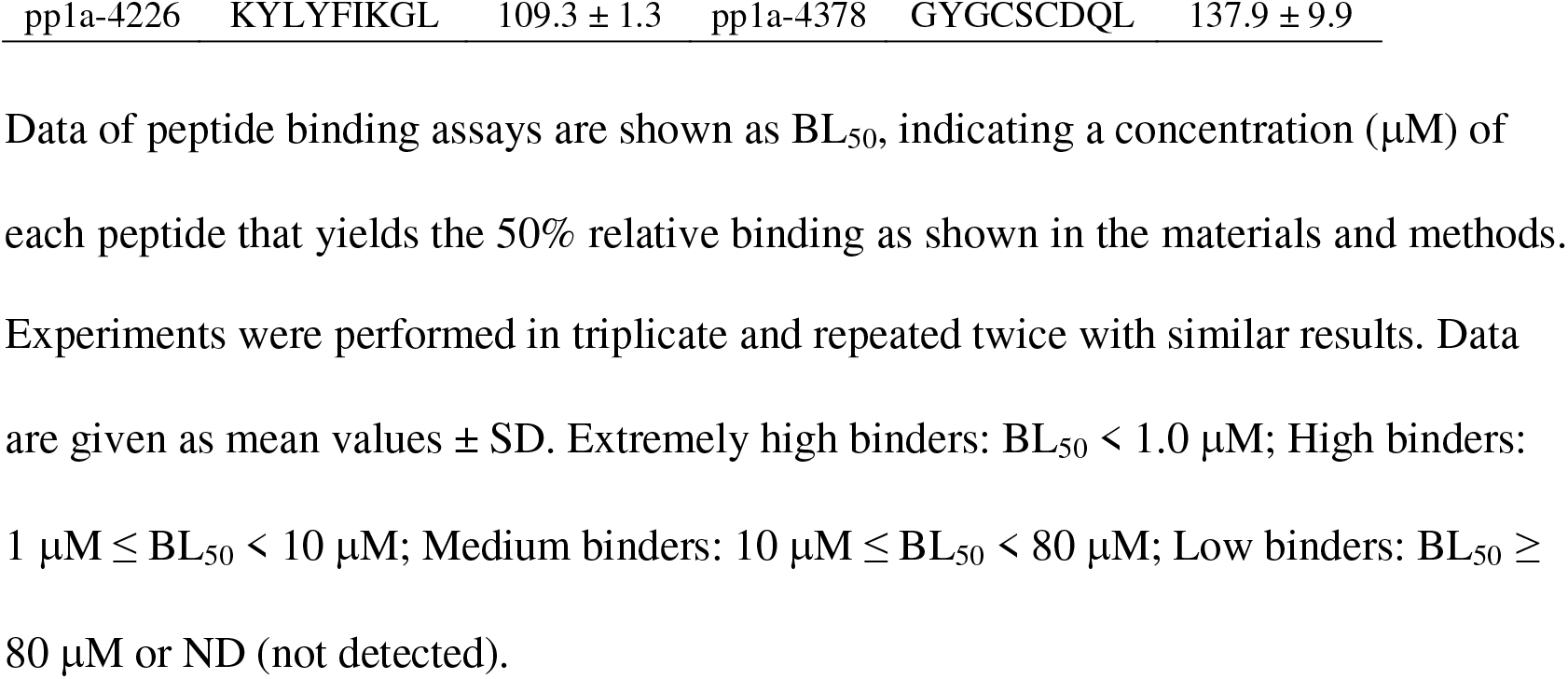
Binding affinities of predicted SARS-CoV-2 pp1a peptides to HLA-A*24:02

**TABLE 3.** Comparison between the peptide binding affinity and the rank^a^ of peptides in the 4 algorithms

|  | Rank | Extremely High | High binder | Medium binder | Low binder |
| --- | --- | --- | --- | --- | --- |
| | | $BL_{50} < 1.0$ (μM) | $1 \leq BL_{50} < 10$ | $10 \leq BL_{50} < 80$ | $BL_{50} \geq 80$ |
| SYFPEITHI | A | 5/80 (6.3%) | 3/80 (3.8%) | 3/80 (3.8%) | 9/80 (11.3%) |
|  | B | 5/80 (6.3%) | 4/80 (5.0%) | 5/80 (6.3%) | 10/80 (10.0%) |
|  | C | 1/80 (1.3%) | 0/80 (0%) | 5/80 (6.3%) | 17/80 (21.3%) |
|  | D | 0/80 (0%) | 3/80 (3.8%) | 2/80 (2.5%) | 8/80 (10.0%) |
| IEDB | A | 9/80 (11.3%) | 4/80 (5.0%) | 2/80 (2.5%) | 7/80 (8.8%) |
|  | B | 2/80 (2.5%) | 4/80 (5.0%) | 8/80 (10.0%) | 13/80 (16.3%) |
|  | C | 0/80 (0%) | 2/80 (2.5%) | 5/80 (6.3%) | 12/80 (15.0%) |
|  | D | 0/80 (0%) | 0/80 (0%) | 0/80 (0%) | 12/80 (15.0%) |
| ProPred-I | A | 4/80 (5.0%) | 3/80 (3.8%) | 4/80 (5.0%) | 9/80 (11.3%) |
|  | B | 7/80 (8.8%) | 2/80 (2.5%) | 3/80 (3.8%) | 8/80 (10.0%) |
|  | C | 0/80 (0%) | 2/80 (2.5%) | 4/80 (5.0%) | 16/80 (20.0%) |
|  | D | 0/80 (0%) | 3/80 (3.8%) | 4/80 (5.0%) | 11/80 (13.8%) |
| NetCTL | A | 9/80 (11.3%) | 1/80 (1.3%) | 0/80 (0%) | 7/80 (8.8%) |
|  | B | 1/80 (1.3%) | 7/80 (8.8%) | 9/80 (11.3%) | 5/80 (6.3%) |
|  | C | 1/80 (1.3%) | 1/80 (1.3%) | 3/80 (3.8%) | 15/80 (18.8%) |
|  | D | 0/80 (0%) | 1/80 (1.3%) | 3/80 (3.8%) | 17/80 (21.3%) |

<sup>a</sup>Rank: Peptides were classified into four ranks (A: Excellent; B: Very good; C: good; D: Poor) in each of the four algorithms (SYFPEITHI, IEDB, ProPred-I, NetCTL).

In the following experiments, 36 peptides involving extremely high, high, and medium binders were chosen to investigate their abilities of peptide-specific CTL induction.

### Induction of SARS-CoV-2 pp1a-specific CD8^+^ T cell responses in HLA-A*24:02 transgenic mice immunized with liposomal peptides

The 36 peptides were randomly divided into 6 groups. Six peptides in each group were mixed and encapsulated into liposomes as described in the materials and methods.

HLA-A*24:02 transgenic mice were then subcutaneously (s.c.) immunized four times at a one-week interval with peptide-encapsulated liposomes together with CpG adjuvant. One week later, spleen cells of immunized mice were prepared, stimulated *in vitro* with a relevant peptide for 5 hours, and stained for their expression of cell-surface CD8 and intracellular interferon-gamma (IFN-γ). As shown in Fig. 3, it was demonstrated that significant numbers of IFN-γ-producing CD8^+^ T cells were detected in mice immunized with 22 liposomal peptides including pp1a-265, -634, -835, -1182, -1255, -1417, -1845, -1899, -2330, -2338, -2350, -2590, -2779, -3104, -3114, -3153, -3249, -3606, -3684, -3752, -3792, and -4229.

**FIG 3.**
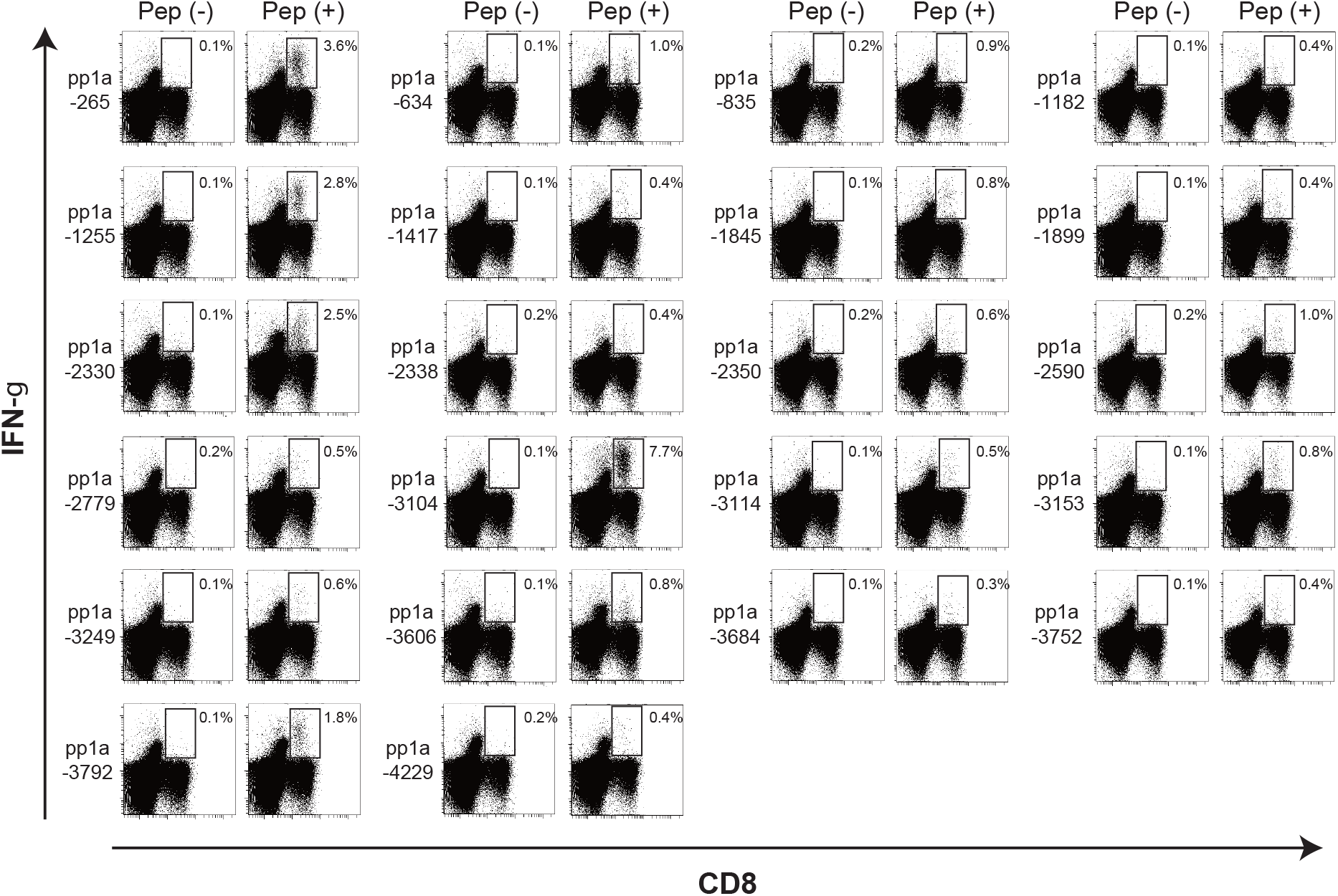
Intracellular IFN-γ staining of CD8^+^ T cells stimulated with peptides derived from SARS-CoV-2 pp1a. After HLA-A*24:02 transgenic mice were immunized with liposomal peptides derived from SARS-CoV-2 pp1a, spleen cells were stimulated with (+) or without (-) a relevant peptide for 5 hours. Cells were stained for their surface expression of CD8 (x axis) and their intracellular expression of IFN-γ (y axis). Numbers shown indicate the percentages of intracellular IFN-γ^+^ cells within CD8^+^ T cells. The data shown are representative of three independent experiments. Three to five mice per group were used in each experiment, and spleen cells of mice per group were pooled.

These data indicated that the 22 peptides were HLA-A*24:02-restricted CTL candidate epitopes derived from SARS-CoV-2 pp1a. However, the induction level of IFN-γ-producing CD8^+^ T cells varied among the 22 peptides. Five peptides including pp1a-265, -1255, -2330, -3104, and -3792 elicited high percentages of intracellular IFN-γ^+^ cells in CD8^+^ T cells, ranging from 1.8% to 7.7%, whereas the other 17 peptides induced medium (0.5-1%) or low percentages (0.1-0.5%) of IFN-γ^+^ CD8^+^ T cells (Fig. 3). When comparing between the data of ICS and the peptide binding affinity (Table 4), it was shown that all of extremely high binders did not elicit IFN-γ producing CD8^+^ T cells and two medium binder peptides activated high percentages of intracellular IFN-γ^+^ cells in CD8^+^ T cells. However, the proportion of extremely high binder peptides that induced IFN-γ producing CD8^+^ T cells was higher than that of medium binder peptides (Table 4), confirming that the peptide binding affinity to HLA class I molecules is closely associated with the induction of peptide-specific CTLs.

**TABLE 4.**
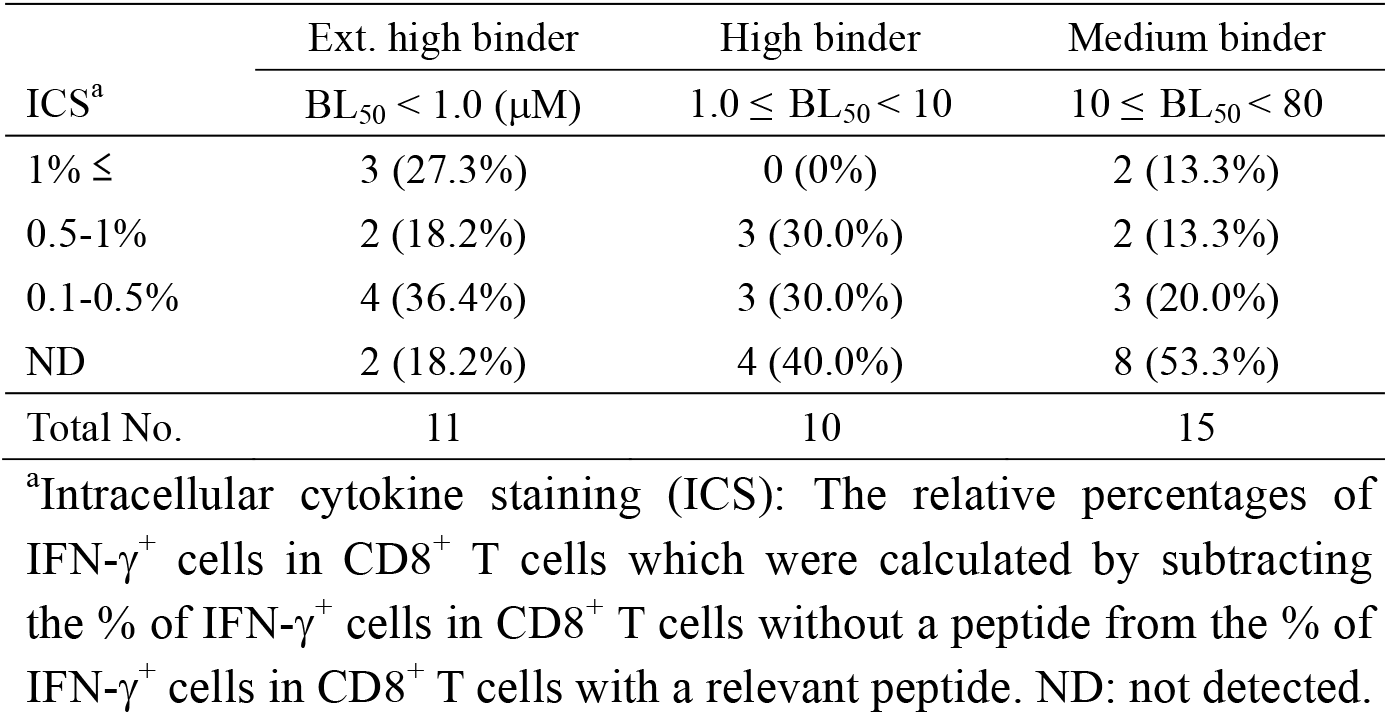
Correlation between the peptide binding affinity and the peptide immunogenicity

### Conservation analysis of CTL epitopes in the database of SARS-CoV-2 variants

We next investigated whether the 22 candidate epitopes were mutated in various SARS-CoV-2 variants. To do this, we utilized the National Center for Biotechnology Information (NCBI) Virus database (https://www.ncbi.nlm.nih.gov) (38), in which they provide us data-sets of mutations in the Sequence Read Archive (SRA) records of SARS-CoV-2 variants. In the database, the nucleotide and amino acid sequences of variants in SRA records were aligned for comparison with those of the original strain, Wuhan-Hu-1 (NCBI Reference Sequence: NC_045512.2). In the SRA mutation data, the most frequent, non-synonymous amino acid change was the mutation from D to G at position 614 (D614G) in the S protein, and the total count of D614G across the database was 615,601 in 924,785 SRA runs (Frequency per run: 66.6%) available as of 23rd August 2021. To investigate the conservation of the 22 epitopes, we counted the total number of non-synonymous amino acid substitutions present in the 9-mer amino acid sequence of each epitope that were found in a number of SRA sequencing data of SARS-CoV-2 variants in 924,785 SRA runs. It was discovered that all of those epitopes had more or less amino acid substitutions in their amino acid sequences (Table 5 & Fig. 4), indicating none of them were fully conserved throughout all of the available SRA data. However, there were seven epitopes with low counts of total mutations present in their 9-mer amino acid sequences, indicating that the amino acid sequences of the seven epitopes were hardly affected by a number of mutations in the SRA database (Table 5). The 7 epitopes were pp1a-835 (559 count in 924,785 SRA runs; Frequency per run: 0.06%), -1417 (245; Frequency: 0.03%), -1899 (531; Frequency: 0.06%), -2590 (611; Frequency: 0.07%), -3104 (142; Frequency: 0.02%), -3792 (336; Frequency: 0.04%) and -4229 (83; Frequency: 0.01%) (Table 5). The number of mutations at each amino acid position in the 9-mer amino acid sequence of an epitope was shown in Fig. 4. In contrast, numbers of amino acid changes in the epitope sequences were very high in some other epitopes including pp1a-265 (53,049; Frequency per run: 5.74%), -2779 (18,819; Frequency: 2.03%), -3249 (126,956; Frequency: 13.73%), and -3606 (19,733; Frequency: 2.13%) (Table 5).

**FIG 4.**
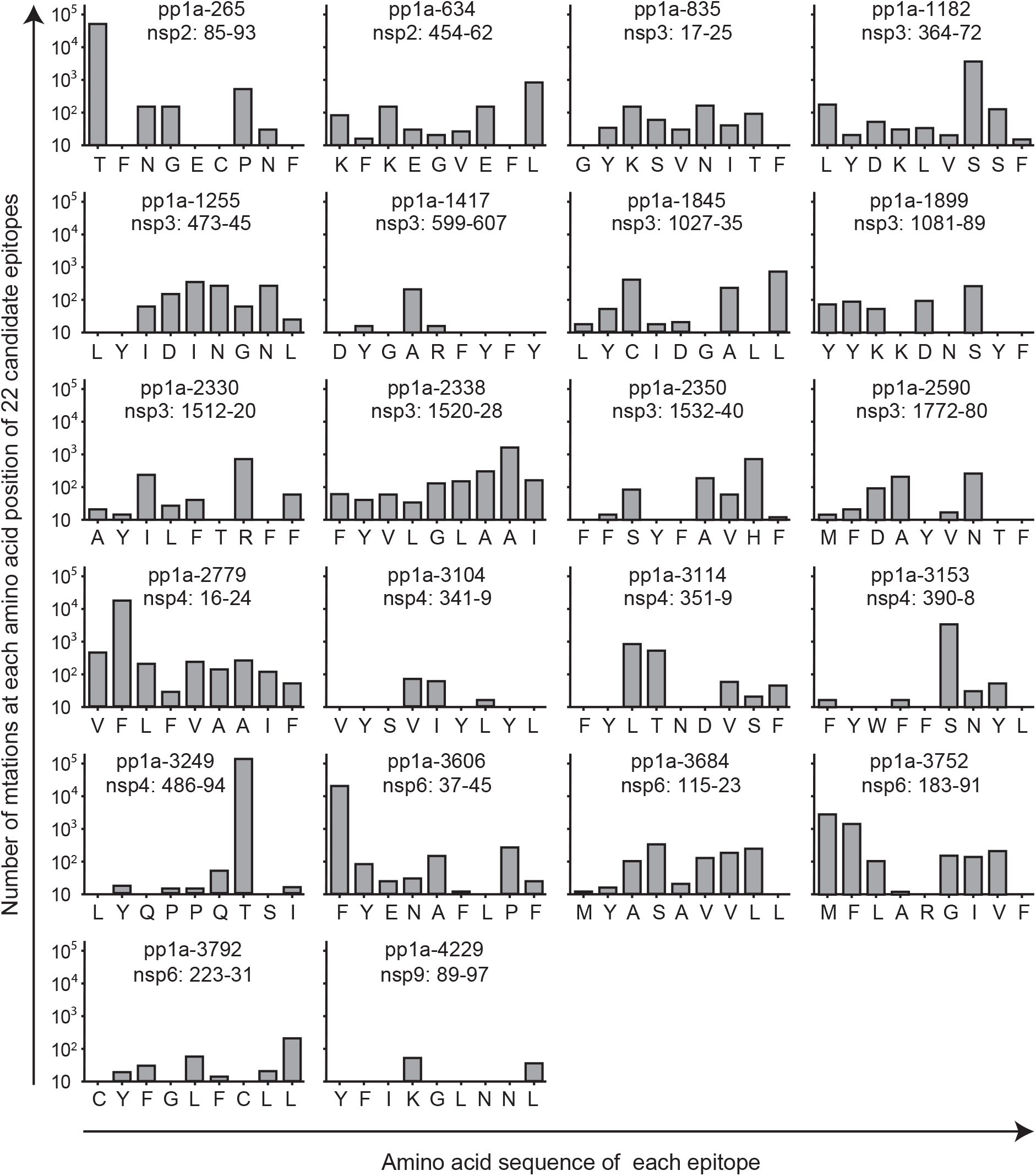
Number of the total non-synonymous mutations at each amino acid position of 22 candidate epitopes. Number of the total non-synonymous amino acid substitutions at each amino acid position of 22 candidate epitopes was counted using the SRA data of SARS-CoV-2 variants in the NCBI Virus database.

**TABLE 5.**
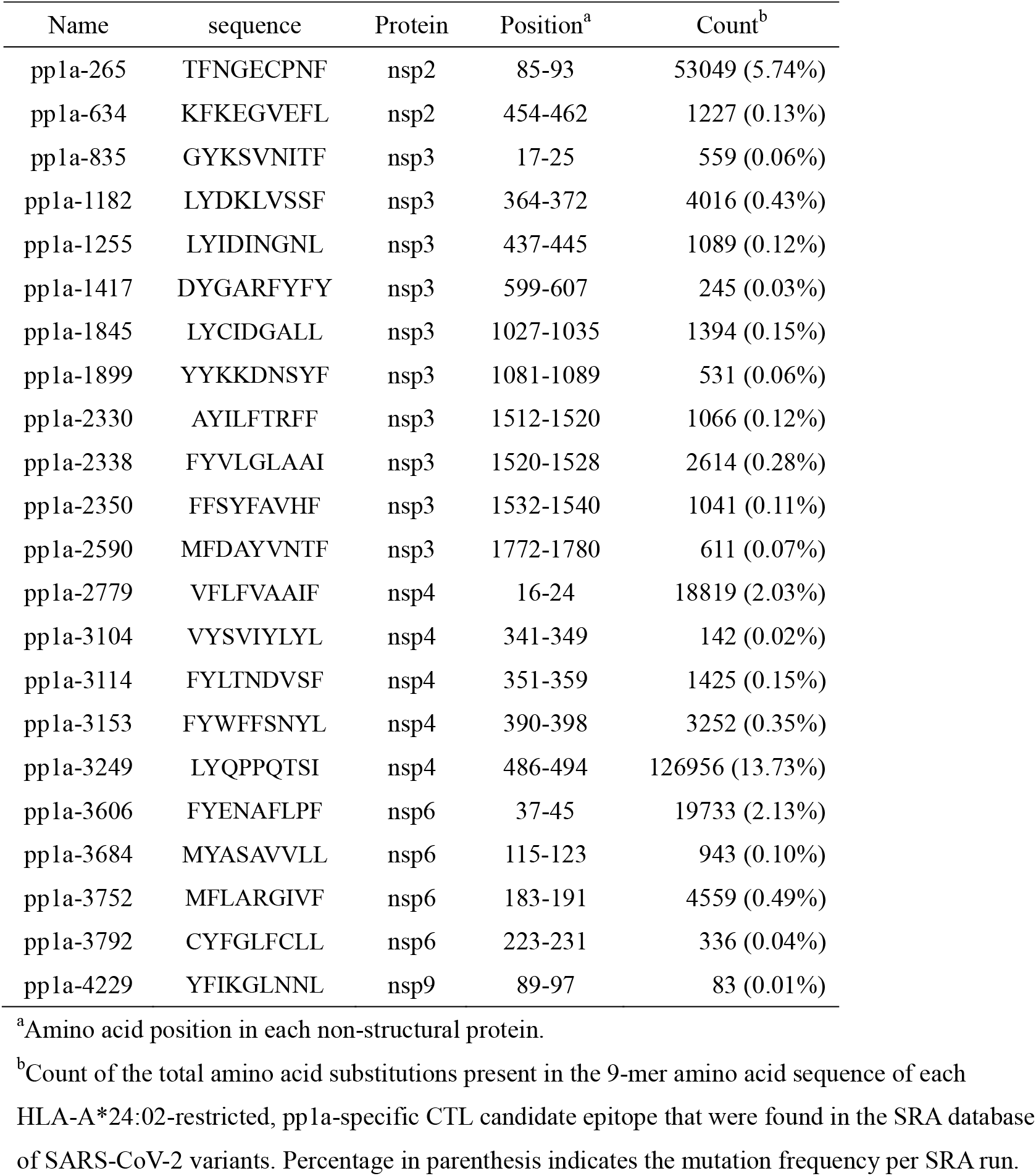
Count of total non-synonymous amino acid changes in each of the 22 HLA-A*24:02-restricted, pp1a-specific CTL candidate epitopes

It was then determined which of the relatively conserved top 4 epitopes, namely pp1a-1417, -3104, -3792, and -4229, was most dominant in the induction of pp1a-specific CTLs. The same amounts of the 4 peptide solutions at an equal concentration were mixed together and encapsulated into liposomes. Eight mice were immunized with the liposomes containing the peptide mixture. One week later, spleen cells were incubated with each of the 4 peptides for 5 hours, and the ICS assay was performed. It was found that pp1a-3104 was far superior to all other peptides in the induction of peptide-specific IFN-γ^+^ CD8^+^ T cells (Fig. 5A). We also examined the peptide-specific induction of CD8^+^ T cells expressing a degranulation marker, CD107a. As shown in Fig. 5B, pp1a-3104 was statistically predominant over pp1a-1417 and -4229 for the CD107a induction of CD8^+^ T cells. Thus, it was found that pp1a-3104 was the most prominent HLA-A*24:02-restricted CTL epitope among the conserved top 4 epitopes.

**FIG 5.**
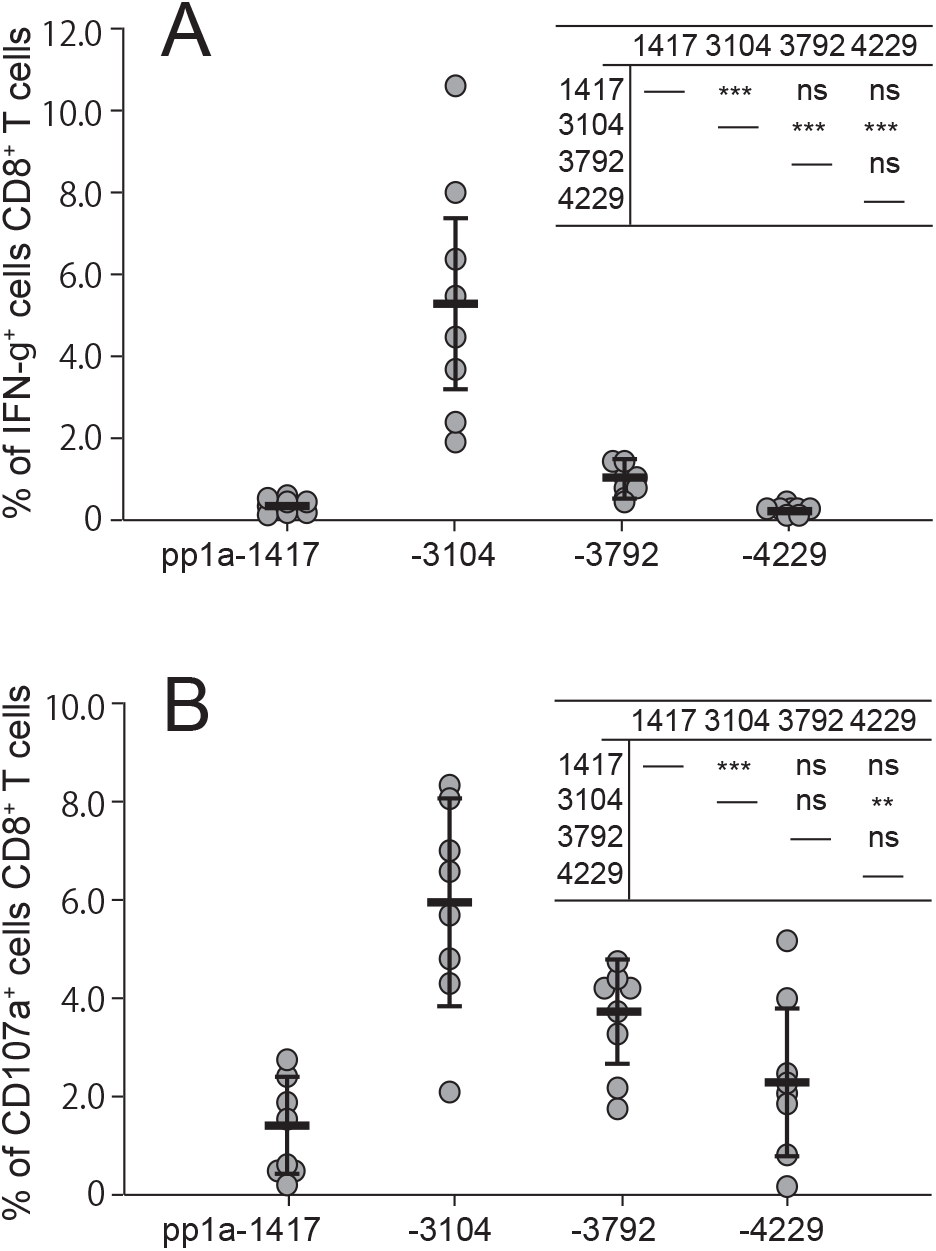
Comparison of the conserved top 4 peptides in the induction of IFN-γ^+^ CD8^+^ T cells (A) and CD107a^+^ CD8^+^ T cells (B). Eight mice were immunized with the mixture of 4 peptides involving pp1a-1417, -3104, -3792, and -4229 in liposomes with CpG. After one week, spleen cells were stimulated with or without each of the 4 peptides, and the expression of intracellular IFN-γ (A) or CD107a (B) in CD8^+^ T cells was stained. Data indicate the relative percentages of IFN-γ^+^ (A) and CD107a^+^ (B) cells in CD8^+^ T cells which were obtained by subtracting the % of IFN-γ^+^ and CD107a^+^ cells in CD8^+^ T cells without a peptide from the % of IFN-γ^+^ and CD107a^+^ cells in CD8^+^ T cells with a peptide, respectively. Each gray circle represents an individual mouse. Data are shown as the mean (horizontal bars) ± SD. Statistical analyses of the data among the 4 peptides in Fig. 5A and Fig. 5B were performed by one-way ANOVA followed by post-hoc tests. Results of statistical analyses were shown as a table in the upper right corner of each figure. **, *P* < 0.01; ***, *P* < 0.001; ns, not significant.

## Discussion

All of the current available COVID-19 vaccines have been directed against the S protein of the original SARS-CoV-2, and therefore they are less effective against some variants with mutated S such as the Beta and Delta strains than the original virus. Our concern is that SARS-CoV-2 is currently under evolution and various variants are appearing one after another. One day soon, new mutant strains that perfectly evade the immunity generated by the vaccines may emerge. To develop a next-generation vaccine to compensate for the viral evolution, it may be beneficial to take advantage of CTLs because they can target a wide range of SARS-CoV-2-derived proteins, involving comparatively conserved non-structural proteins.

Here, we have identified twenty-two HLA-A*24:02-restricted CTL candidate epitopes derived from SARS-CoV-2 pp1a using HLA-A*24:02 transgenic mice. The pp1a is a large polyprotein consisting of 4,401 amino acids that may be relatively conserved compared to structural proteins such as the S protein (31). Furthermore, Tarke et al. demonstrated that most of T cell epitopes they identified were conserved across the Alpha, Beta, Gamma, and Epsilon (CAL.20C) variants, and the impact of the four variants on the total CD8^+^ T cell reactivity in vaccinated individuals was negligible (39). Hence, we firstly thought it might be possible to find pp1a-derived epitopes that were fully conserved across a number of the existing SARS-CoV-2 variants. Unfortunately, none of the 22 epitopes we identified were found to be completely conserved throughout vast amounts of the SRA data in the NCBI Virus database. This is understandable because most (73.3%) of the 4,401 amino acids that make up the pp1a have non-synonymous amino acid substitutions found in the SARS-CoV-2 SRA data. As shown in Table 5, however, seven epitopes including pp1a-835, -1417, -1899, -2590, -3104, -3792, and -4299 were relatively conserved due to low counts of total mutations and minimum mutation frequencies of less than 0.1% in their amino acid sequences. Of note, pp1a-3104 was indicated to be the most dominant epitope in the induction of activated CD8^+^ T cells (Fig. 5).

In the current study, we have focused on HLA-A*24:02-restricted CTL epitopes because HLA-A*24:02 is predominant in East Asian people (34, 40) such as Japanese (allele frequency: 32.7%) (41). On the other hand, HLA-A*02:01 individuals are well known to be highly frequent all over the world (34). We previously identified eighteen of HLA-A*02:01-restricted CTL candidate epitopes derived from SARS-CoV-2 pp1a using HLA-A*02:01 transgenic mice (42). Then, we here examined how much these epitopes were mutated across the vast SRA data. As shown in Table 6, four epitopes involving pp1a-2785, -2884, -3403, and -3583 were found to be relatively conserved because of their low mutation frequencies per SRA run. Fig. 6 indicates where the four HLA-A*02:01-restricted (Table 6), and seven HLA-A*24:02-rescricted (Table 5) epitopes with minimum mutation frequencies of less than 0.1% are located in the pp1a, indicating that these epitopes are interspersed in the five non-structural proteins. If the nucleotide sequences encoding some of these CTL epitopes are inserted into the current mRNA vaccine or adenoviral-vectored vaccine, the new vaccine would be effective against almost all of the existing and presumably upcoming variants in HLA-A*02:01 and/or A*24:02 positive individuals who are equivalent to a significant proportion of the world’s population. The new vaccine would elicit both virus-neutralizing antibodies directed against the S protein and pp1a-derived conserved epitope-specific CTLs targeting cells infected with most of the variants.

**FIG 6.**
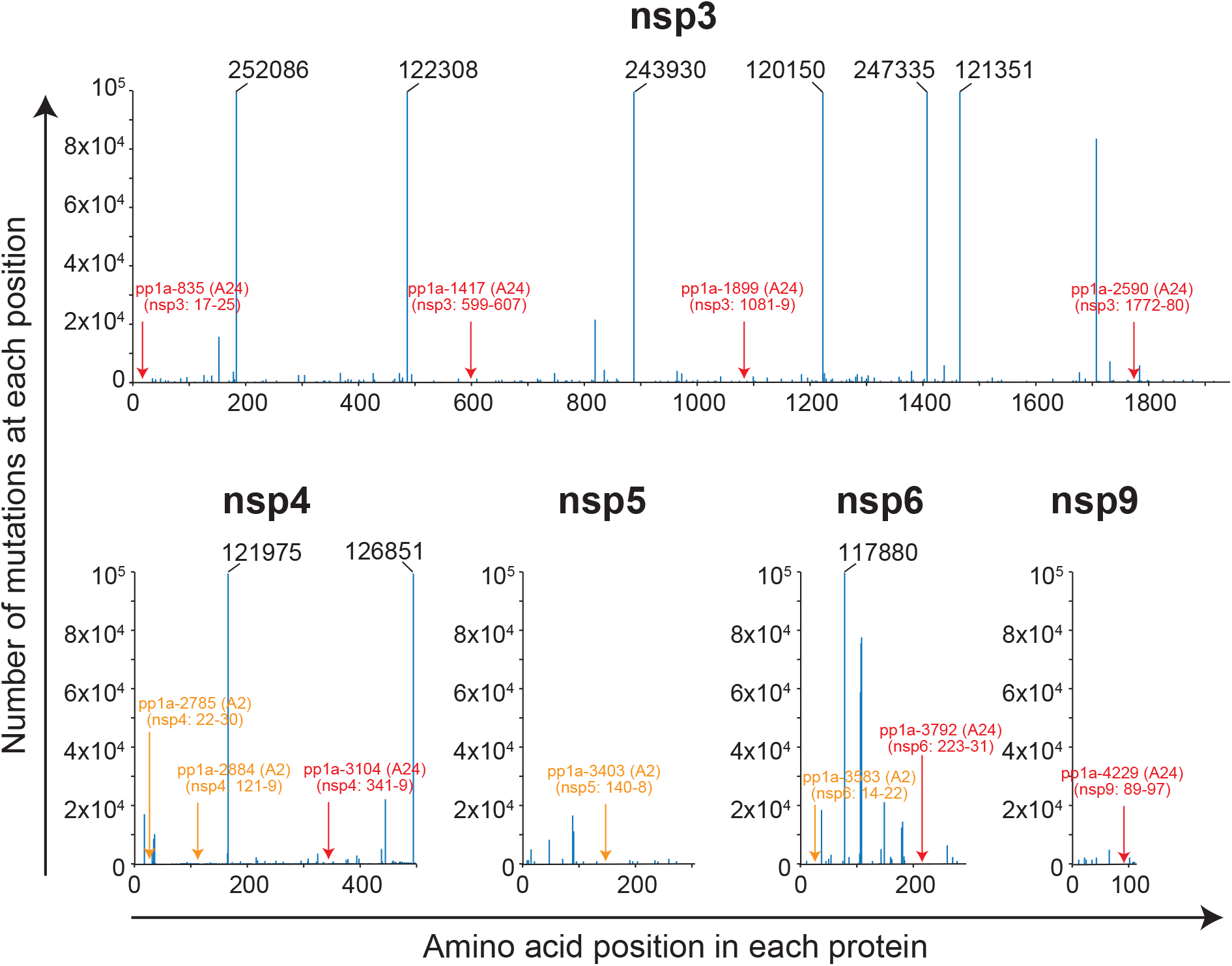
Locations of conserved CTL epitopes in the pp1a. Seven HLA-A*24:02-rescricted (red letters and arrows) and four HLA-A*02:01-restricted (orange letters and arrows) epitopes were selected as conserved epitopes because they demonstrated low mutation frequencies per SRA run of less than 0.1% (Tables 5 & 6). Locations of the eleven conserved CTL epitopes were shown in this figure. The blue line indicates the number of total non-synonymous amino acid substitutions at each amino acid position that were found in a number of SRA sequencing data of SARS-CoV-2 variants. When the number exceeds 10^5^, the actual number is shown at the top of the blue line.

**TABLEL 6.**
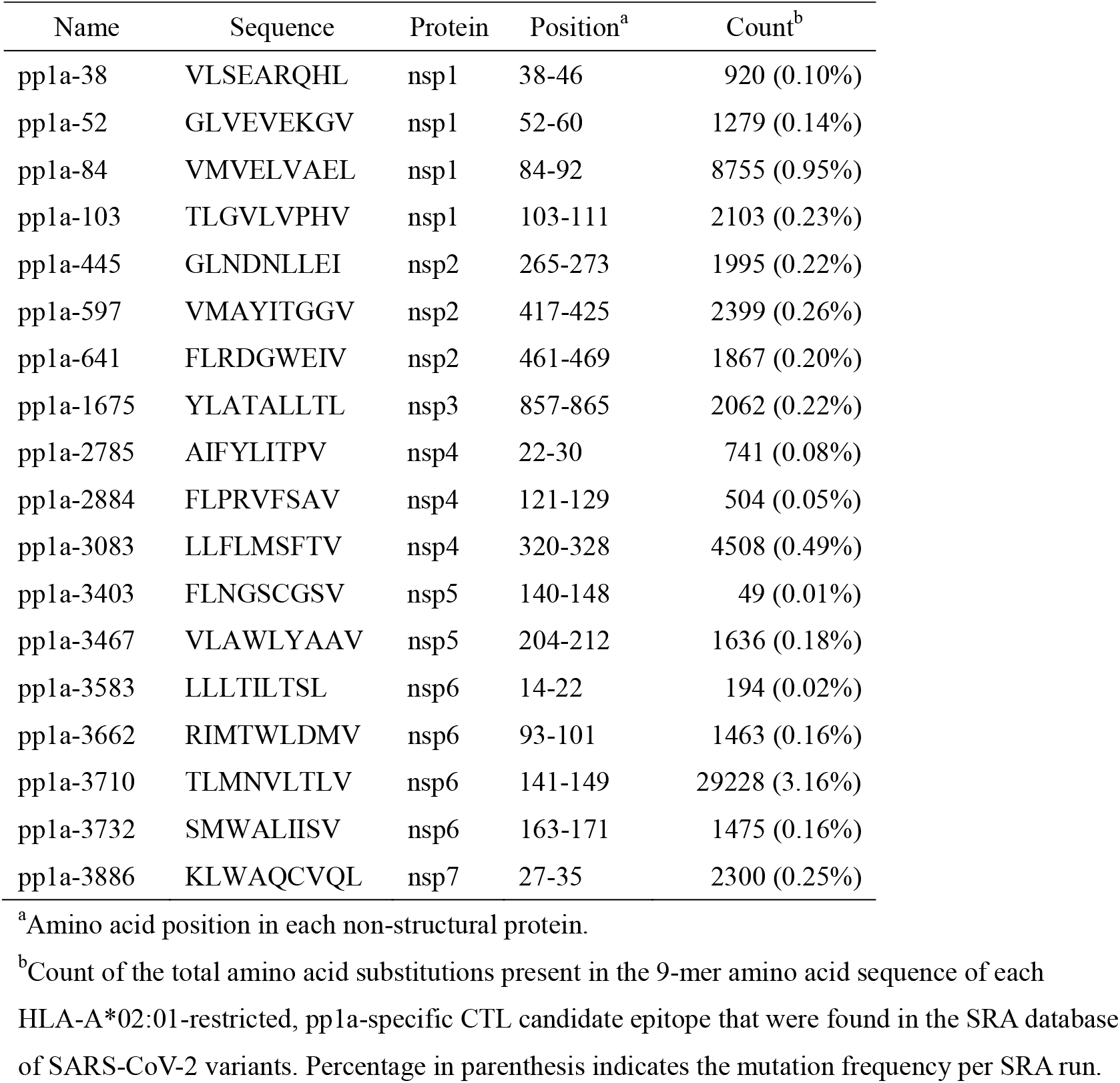
Count of total non-synonymous amino acid changes in each of the 18 HLA-A*02:01-restricted, pp1a-specific CTL candidate epitopes identified in the previous study (42)

To identify HLA-A*24:02-restricted CTL epitopes, we utilized highly reactive HLA-A*24:02 transgenic mice (43). One reason for using MHC-I transgenic mice instead of lymphocytes of SARS-CoV-2-infected individuals is that a large number of lymphocytes are required to examine many candidates of CTL epitopes. Furthermore, when using patients’ lymphocytes, we are only testing whether the peptide candidates are recognized by memory CTLs. In contrast, naive mice can be used to see if the epitope candidates are able to prime peptide-specific CTLs. This may be a better criterion to judge them as vaccine antigens. However, we have to take into account that the immunogenic variation in HLA class I transgenic mice may not be identical to that in humans because the antigen processing and presentation differ between them. In addition, we did not present data showing that viral infection in a mouse model induces T cells targeting these epitopes because liposomal peptides were used as an immunogen. Hence, there is no guarantee that the candidate epitopes identified here are real pp1a-derived epitopes that are presented by human cells during live infection with SARS-CoV-2. Recently, eight epitopes with the same amino acid sequences as pp1a-265, -1182, -1899, -2330, -3104, -3114, -3249, and -3684 (Table 5) have been submitted to the Virus Pathogen Database and Analysis Resource as HLA-A*24:02-restricted pp1a-specific CTL epitopes. Five (pp1a-265, -1182, -1899, -2330, and -3249) of them were shown to be positive in T cell assays using human lymphocytes, and therefore, they are thought to be real epitopes. Three epitopes (pp1a-3104, -3114, and -3684) of them were positive in the binding assay but negative in T cell assays, suggesting that they are not likely to be real epitopes. Then, the remaining 14 candidate epitopes in Table 5 represent new candidate epitopes that have not been previously identified.

In summary, we have identified 22 kinds of HLA-A*24:02-restricted CTL candidate epitopes derived from the pp1a of SARS-CoV-2 using computational algorithms, HLA-A*24:02 transgenic mice and the peptide-encapsulated liposomes. The conservation analysis revealed that the amino acid sequences of 7 out of the 22 epitopes were hardly affected by a number of mutations in the SRA database of SARS-CoV-2 variants. We also found four relatively conserved epitopes among 18 HLA-A*02:01-restricted CTL candidate epitopes that we had previously identified. The new mRNA or adenoviral-vectored vaccine containing nucleotide sequences encoding some of these epitopes might have the potential to become the universal vaccine against almost all of the existing and upcoming SARS-CoV-2 variants.

## Materials and Methods

### Prediction of HLA-A*2402-restricted CTL epitopes

A T-cell epitope database, SYFPEITHI (33) was used to predict HLA-A*24:02-restricted CTL epitopes derived from pp1a of SARS-CoV-2 (GenBank accession numbers: LC528232.1 & LC528233.1). Eighty of 9-mer peptides with superior scores (17 or higher) in the SYFPEITHI database were selected (Table 1) and were synthesized by Eurofins Genomics (Tokyo, Japan). These epitopes were also evaluated by three other algorithms, IEDB (34), ProPred-1 (35), and NetCTL (36) (Table 1). An HLA-A*24:02-restricted control peptide, Influenza PA_130-138_ (sequence: YYLEKANKI) (37), was synthesized as well.

### Mice

We used HLA-A*24:02 transgenic mice which were kindly provided by Dr. François A. Lemonnier (Pasteur Institute, Paris, France). The HLA-A*24:02 transgenic mouse expresses an HLA-A*24:02 monochain, designated as HHD-A24, in which human β2m is covalently linked to a chimeric heavy chain composed of HLA-A*24:02 (α1 and α2 domains) and H-2D^b^ (α3, transmembrane, and cytoplasmic domains) in an H-2D^b^, K^b^, and mouse β2m triple knockout environment (43). Six- to ten-week-old mice were used for all experiments. Mice were housed in appropriate animal care facilities at Saitama Medical University, and were handled according to the international guideline for experiments with animals. This study was approved by the Animal Research Committee of Saitama Medical University.

### Cell line

The HHD-A24 gene, which was composed of human β2m cDNA linked to the chimeric heavy chain cDNA encoding α1/α2 domains of HLA-A*24:02, and α3/transmembrane/cytoplasmic domains of H-2D^b^, was synthesized by Eurofins Genomics.

HHD-A24 cDNA was subcloned into the mammalian expression plasmid, pcDNA3.1 (+) (Thermo Fisher Scientific, MA) (pcDNA3.1-HHD-A24). The TAP2-dificient mouse lymphoma cell line, RMA-S (H-2^b^) was transfected with pcDNA3.1-HHD-A24 by electroporation (Gene Pulser Xcell, Bio-Rad, Hercules, CA), and cloned by the FACSAria II cell sorter (BD Biosciences, Franklin Lakes, NJ). The resultant RMA-S-HHDA-24 cell line was cultured in RPMI-1640 medium (Nacalai Tesque Inc., Kyoto, Japan) with 10% FCS (Biowest, Nuaille, France) and 500 μg/ml G418 (Nacalai Tesque Inc.)

### Peptide binding assay

Binding affinity of each peptide to HLA-A*24:02 was measured by the peptide binding assay using RMA-S-HHD-A24 cells, as described before (42). In brief, RMA-S-HHD-A24 cells were pre-cultured overnight at 26°C in a CO_2_ incubator, and pulsed with each peptide at various concentrations for 1 hour at 26°C. Peptide-pulsed cells were incubated for 3 hours at 37°C, and were stained with anti-HLA-A24 monoclonal antibody (mAb), A11.1M (44), followed by FITC-labeled goat anti-mouse IgG antibody (Sigma-Aldrich, St. Louis, MO).

Mean fluorescence intensity (MFI) of HLA-A*24:02 expression on the surface of RMA-S-HHD-A24 cells was measured by flow cytometry (FACSCanto II, BD Biosciences), and standardized as the percent cell surface expression by the following formula: % relative binding = [{(MFI of cells pulsed with each peptide) – (MFI of cells incubated at 37°C without a peptide)}/{(MFI of cells incubated at 26°C without a peptide) – (MFI of cells incubated at 37°C without a peptide)}] × 100. The concentration of each peptide that yields the 50% relative binding was calculated as the half-maximal binding level (BL_50_).

### Peptide-encapsulated liposomes

Peptide-encapsulated liposomes were prepared using Lipocapsulater FD-U PL (Hygieia BioScience, Osaka, Japan), as previously described (42). Briefly, each of synthetic peptides was dissolved in DMSO at a final concentration of 10 mM. For the first screening of HLA-A*24:02-restricted epitopes, 20 μl each of 6 peptide solutions was mixed together, and the total volume was increased to 2 ml by adding H_2_O. For the identification of dominant epitopes, 20 μl each of 10 mM peptides selected was mixed together, and diluted to 2 ml with H_2_O. The peptide solution was added into a vial of Lipocapsulater containing 10 mg of dried liposomes, and incubated for 15 min at room temperature. The resultant solution contains peptide-encapsulated liposomes.

### Immunization

Mice were immunized s.c. four times at a one-week interval with peptide-encapsulated liposomes (100 μl/mouse for priming and 50 μl/mouse for boosting) together with CpG-ODN (5002: 5’-TCCATGACGTTCTTGATGTT-3’, Hokkaido System Science, Sapporo, Japan) (5 μg/mouse) in the footpad.

### Intracellular cytokine staining (ICS)

ICS was performed as described previously (42). Spleen cells of immunized mice were incubated with 50 μM of each peptide for 5 hours at 37°C in the presence of brefeldin A (GolgiPlug, BD Biosciences), and were stained with FITC-conjugated anti-mouse CD8 mAb (BioLegend, San Diego, CA). Cells were then fixed, permeabilized, and stained with phycoerythrin (PE)-conjugated rat anti-mouse IFN-γ mAb (BD Biosciences). After washing the cells, flow cytometric analyses were performed using flow cytometry (FACSCanto II, BD Biosciences).

### Conservation analysis of CTL epitopes

To examine the conservation of the CTL candidate epitopes, we utilized the SRA data of SARS-CoV-2 variants in the NCBI Virus database. We counted the total number of non-synonymous amino acid changes present in the 9-mer amino acid sequence of each epitope that were found in the SRA mutation database, and calculated percentage of the mutation frequency per SRA run of each epitope.

### Detection of CD107a molecules on CD8^+^ T cells

For the detection of CD107a, spleen cells of immunized mice were incubated with 50 μM of each peptide for 6 hours at 37°C in the presence of monensin (GolgiStop, BD Biosciences) and 0.8 μg of FITC-conjugated anti-mouse CD107a mAb (BioLegend). Cells were then stained with PE-Cy5-conjugated anti-mouse CD8 mAb (BioLegend), and were analyzed by flow cytometry (FACSCanto II, BD Biosciences).

### Statistical analyses

One-way ANOVA followed by post-hoc tests was performed for statistical analyses among multiple groups using Graphpad Prism 5 software (GraphPad software, San Diego, CA). A value of *P* < 0.05 was considered statistically significant.

## ACKNOWLEDGMENTS

The authors are grateful to Professor François A. Lemonnier (Pasteur Institute, Paris, France) for providing HLA-A*24:02 transgenic mice. This work was supported by a Grant-in-Aid for Scientific Research (C) (JSPS KAKENHI Grant Number: JP18K06631) to M. M., and a Grant-in-Aid for Early-Career Scientists (JSPS KAKENHI Grant Number: JP18K15430) to A. T. from Japan Society for the Promotion of Science. The authors have no conflicting financial interests.

## References

1. Polack FP, Thomas SJ, Kitchin N, Absalon J, Gurtman A, Lockhart S, Perez JL, Marc GP, Moreira ED, Zerbini C, Bailey R, Swanson KA, Roychoudhury S, Koury K, Li P, Kalina WV, Cooper D, Frenck Jr. RW, Hammitt LL, Türeci O, Nell H, Schaefer A, Unal S, Tresnan DB, Mather S, Dormitzer PR, Şahin U, Jansen KU, Gruber WC, for the C4591001 Clinical Trial Group. 2020. Safety and efficacy of the BNT162b2 mRNA Covid-19 vaccine. N Engl J Med 383:2603–2615.

2. Baden LR, El Sahly HM, Essink B, Kotloff K, Frey S, Novak R, Diemert D, Spector SA, Rouphael N, Creech CB, McGettigan J, Khetan S, Segall N, Solis J, Brosz A, Fierro C, Schwartz H, Neuzil K, Corey L, Gilbert P, Janes H, Follmann D, Marovich M, Mascola J, Polakowski L, Ledgerwood J, Graham BS, Bennett H, Pajon R, Knightly C, Leav B, Deng W, Zhou H, Han S, Ivarsson M, Miller J, Zaks T, for the COVE Study Group. 2021. Efficacy and safety of the mRNA-1273 SARS-CoV-2 vaccine. N Engl J Med 384:403–416.

3. Bok K, Sitar S, Graham BS, Mascola JR. 2021. Accelerated COVID-19 vaccine development: milestones, lessons, and prospects. Immunity 54:1636–1651.

4. Teijaro JR, Farber DL. 2021. COVID-19 vaccines: modes of immune activation and future challenges. Nat Rev Immunol 21:195–197.

5. Robson F, Khan KS, Le TK, Paris C, Demirbag S, Barfuss P, Rocchi P, Ng WL. 2020. Coronavirus RNA proofreading: Molecular basis and therapeutic targeting. Mol Cell 79:710–727.

6. Muik A, Wallisch A-K, Sänger B, Swanson KA, Mühl J, Chen W, Cai H, Maurus D, Sarkar R, Türeci Ö, Dormitzer PR, Sahin U. 2021. Neutralization of SARS-CoV-2 lineage B.1.1.7 pseudovirus by BNT162b2 vaccine–elicited human sera. Science 371:1152–1153.

7. Planas D, Bruel T, Grzelak L, Guivel-Benhassine F, Staropoli I, Porrot F, Planchais C, Buchrieser J, Rajah MM, Bishop E, Albert M, Donati F, Prot M, Behillil S, Enouf V, Maquart M, Smati-Lafarge M, Varon E, Schortgen F, Yahyaoui L, Gonzalez M, De Sèze J, Péré H, Veyer D, Sève A, Simon-Lorière E, Fafi-Kremer S, Stefic K, Mouquet H, Hocqueloux L, van der Werf S, Prazuck T, Schwartz O. 2021. Sensitivity of infectious SARS-CoV-2 B.1.1.7 and B.1.351 variants to neutralizing antibodies. Nat Med 27:917–924.

8. Wang R, Zhang Q, Ge J, Ren W, Zhang R, Lan J, Ju B, Su B, Yu F, Chen P, Liao H, Feng Y, Li X, Shi X, Zhang Z, Zhang F, Ding Q, Zhang T, Wang X, Zhang L. 2021. Analysis of SARS-CoV-2 variant mutations reveals neutralization escape mechanisms and the ability to use ACE2 receptors from additional species. Immunity 54:1611–1621.

9. Alter G, Yu J, Liu J, Chandrashekar A, Borducchi EN, Tostanoski LH, McMahan K, Jacob-Dolan C, Martinez DR, Chang A, Anioke T, Lifton M, Nkolola J, Stephenson KE, Atyeo C, Shin S, Fields P, Kaplan I, Robins H, Amanat F, Krammer F, Baric RS, Gars ML, Sadoff J, de Groot AM, Heerwegh D, Struyf F, Douoguih M, van Hoof J, Schuitemaker H, Barouch DH. 2021. Immunogenicity of Ad26.COV2.S vaccine against SARS-CoV-2 variants in humans. Nature 596:268–272.

10. Cele S, Gazy I, Jackson L, Hwa S-H, Tegally H, Lustig G, Giandhari J, Pillay S, Wilkinson E, Naidoo Y, Karim F, Ganga Y, Khan K, Bernstein M, Balazs AB, Gosnell BI, Hanekom W, Moosa M-YS, Network for Genomic Surveillance in South Africa, COMMIT-KZN Team, Lessells RJ, de Oliveira T, Sigal A. 2021. Escape of SARS-CoV-2 501Y.V2 from neutralization by convalescent plasma. Nature 593:142–146.

11. Chen RE, Zhang X, Case JB, Winkler ES, Liu Y, VanBlargan LA, Liu J, Errico JM, Xie X, Suryadevara N, Gilchuk P, Zost SJ, Tahan S, Droit L, Turner JS, Kim W, Schmitz AJ, Thapa M, Wang D, Boon ACM, Presti RM, O’Halloran JA, Kim AHJ, Deepak P, Pinto D, Fremont DH, Crowe Jr JE, Corti D, Virgin HW, Ellebedy AH, Shi P-Y, Diamond MS. 2021. Resistance of SARS-CoV-2 variants to neutralization by monoclonal and serum-derived polyclonal antibodies. Nat Med 27:717–726.

12. Wang P, Nair MS, Liu L, Iketani S, Luo Y, Guo Y, Wang M, Yu J, Zhang B, Kwong PD, Graham BS, Mascola JR, Chang JY, 1,5, Yin MT, Sobieszczyk M, Kyratsous CA, Shapiro L, Sheng Z, Huang Y, Ho DH. 2021. Antibody resistance of SARS-CoV-2 variants B.1.351 and B.1.1.7. Nature 593:130–135.

13. Wibmer CK, Ayres F, Hermanus T, Madzivhandila M, Kgagudi P, Oosthuysen B, Lambson BE, de Oliveira T, Vermeulen M, van der Berg K, Rossouw T, Boswell M, Ueckermann V, Meiring S, von Gottberg A, Cohen C, Morris L, Bhiman JN, Moore PL. 2021. SARS-CoV-2 501Y.V2 escapes neutralization by South African COVID-19 donor plasma. Nat Med 27:622–625.

14. Abu-Raddad LJ, Butt AA. 2021. Effectiveness of the BNT162b2 Covid-19 Vaccine against the B.1.1.7 and B.1.351 Variants. N Engl J Med 385:187–189.

15. Chemaitelly H, Yassine HM, Benslimane FM, Al Khatib HA, Tang P, Hasan MR, Malek JA, Coyle P, Ayoub HH, Al Kanaani Z, Al Kuwari E, Jeremijenko A, Kaleeckal AH, Latif AN, Shaik RM, Rahim HFA, Nasrallah GK, Al Kuwari MG, Al Romaihi HE, Al-Thani MH, Al Kha A, Butt AA, Bertollini R, Abu-Raddad LJ. 2021. mRNA-1273 COVID-19 vaccine effectiveness against the B.1.1.7 and B.1.351 variants and severe COVID-19 disease in Qatar. Nat Med 27:1614–1621.

16. 16. Dagpunar J. 2021. Interim estimates of increased transmissibility, growth rate, and reproduction number of the Covid-19 B.1.617.2 variant of concern in the United Kingdom. medRxiv https://doi.org/10.1101/2021.06.03.21258293.

17. Campbell F, Archer. B, Laurenson-Schafer H, Jinnai Y, Konings F, Batra N, Pavlin B, Vandemaele K, Van Kerkhove MD, Jombart T, Morgan O, le Polain de Waroux O. 2021. Increased transmissibility and global spread of SARS-CoV-2 variants of concern as at June Euro Surveill. 26:2100509. doi:10.2807/1560-7917.ES.2021.26.24.2100509.

18. 18. Li B, Deng A, Li K, Hu Y, Li Z, Xiong Q, Liu Z, Guo Q, Zou L, Zhang H, Zhang M, Ouyang F, Juan Su J, Su W, Xu J, Lin H, Sun J, Peng J, Jiang H, Zhou P, Hu T, Luo M, Zhang Y, Zheng H, Xiao J, Liu T, Che R, Zeng H, Zheng Z, Huang Y, Yu J, Yi L, Wu J, Chen J, Zhong H, Deng X, Kang M, Pybus OG, Hall M, Lythgoe KA, Li Y, Yuan J, He J, Lu J. 2021. Viral infection and transmission in a large, well-traced outbreak caused by the SARS-CoV-2 Delta variant. medRxiv https://doi.org/10.1101/2021.07.07.21260122.

19. 19. Fisman DN, Tuite AR. 2021. Progressive Increase in Virulence of Novel SARS-1 CoV-2 Variants in Ontario, Canada. medRxiv https://doi.org/10.1101/2021.07.05.21260050.

20. Bernal JL, Andrews N, Gower C, Gallagher E, Simmons R, Thelwall S, Stowe J, Tessier E, Groves N, Dabrera G, Richard Myers R, Campbell CNJ, Amirthalingam G, Edmunds M, Zambon M, Brown KE, Hopkins S, Chand M, Ramsay M. 2021. Effectiveness of Covid-19 vaccines against the B.1.617.2 (Delta) variant. N Engl J Med 385:585–594.

21. 21. Riemersma KK, Grogan BE, Kita-Yarbro A, Halfmann P, Kocharian A, Florek KR, Westergaard R, Bateman A, Jeppson GE, Kawaoka Y, O’Connor DH, Friedrich TC, Grande KM. 2021. Shedding of infectious SARS-CoV-2 despite vaccination when the Delta variant is prevalent - Wisconsin, July 2021. medRxiv https://doi.org/10.1101/2021.07.31.21261387.

22. Moderbacher CR, Ramirez SI, Dan JM, Grifoni A, Hastie KM, Weiskopf D, Belanger S, Abbott RK, Kim C, Choi J, Kato Y, Crotty EG, Kim C, Rawlings SA, Mateus J, Tse LPV, Frazier A, Baric R, Peters B, Greenbaum J, Saphire EO, Smith DM, Sette A, Crotty S. 2020. Antigen-specific adaptive immunity to SARS-CoV-2 in acute COVID-19 and associations with age and disease severity. Cell 183:996–1012.

23. Peng Y, Mentzer AJ, Liu G, Yao X, Yin Z, Dong D, Dejnirattisai W, Rostron T, Supasa P, Liu C, López-Camacho C, Slon-Campos J, Zhao Y, Stuart DI, Paesen GC, Grimes JM, Antson AA, Bayfield OW, Hawkins DEDP, Ker D-S, Wang B, Turtle L, Subramaniam K, Thomson P, Zhang P, Dold C, Ratcliff J, Simmonds P, de Silva T, Sopp P, Wellington D, Rajapaksa U, Chen Y-L, Mariolina Salio M, Napolitani G, Paes W, Borrow P, Kessler BM, Fry JW, Schwabe NF, Semple MG, Baillie JK, Moore SC, Openshaw PJM, Ansari MA, Dunachie S, Barnes E, Frater J, Kerr G, Goulder P, Lockett T, Levin R, Zhang Y, Jing R, Ho L-P, Oxford Immunology Network Covid-19 Response T cell Consortium, ISARIC4C Investigators, Cornall RJ, Conlon CP, Klenerman P, Screaton GR, Mongkolsapaya J, McMichael A, Knight JC, Ogg G, Dong T. 2020. Broad and strong memory CD4+ and CD8+ T cells induced by SARS-CoV-2 in UK convalescent individuals following COVID-19. Nat Immunol 21:1336–1345.

24. Diao B, Wang C, Tan Y, Chen X, Liu Y, Ning L, Chen L, Li M, Liu Y, Wang G, Yuan Z, Feng Z, Zhang Y, Wu Y, Chen Y. 2020. Reduction and functional exhaustion of T cells in patients with Coronavirus Disease 2019 (COVID-19). Front Immunol 11:827. https://doi.org/10.3389/fimmu.2020.00827.

25. Tan AT, Linster M, Tan CW, Le Bert N, Chia WN, Kunasegaran K, Zhuang Y, Tham CYL, Chia A, Smith GJD, Young B, Kalimuddin S, Low JGH, Lye D, Wang L-F, Bertoletti A. 2021. Early induction of functional SARS-CoV-2-specific T cells associates with rapid viral clearance and mild disease in COVID-19 patients. Cell Rep 34:108728. https://doi.org/10.1016/j.celrep.2021.108728.

26. Soresina A, Moratto D, Chiarini M, Paolillo C, Baresi G, Focà E, Michela Bezzi M, Baronio B, Giacomelli M, Badolato R. 2020. Two X-linked agammaglobulinemia patients develop pneumonia as COVID-19 manifestation but recover. Pediatr Allergy Immunol 31:565–569.

27. McMahan K, Yu J, Mercado NB, Loos C, Tostanoski LH, Chandrashekar A, Liu J, Peter L, Atyeo C, Zhu A, Bondzie EA, Dagotto G, Gebre MS, Jacob-Dolan C, Li Z, Nampanya F, Patel S, Pessaint L, Van Ry A, Blade K, Yalley-Ogunro J, Cabus M, Brown R, Cook A, Teow E, Andersen H, Lewis MG, Lauffenburger DA, Alter G, Barouch DH. 2021. Correlates of protection against SARS-CoV-2 in rhesus macaques. Nature 590:630–634.

28. Arunachalam PS, Scott MKD, Hagan T, Li C, Feng Y, Wimmers F, Grigoryan L, Trisal M, Edara VV, Lai L, Chang SE, Feng A, Dhingra S, Shah M, Lee AS, Chinthrajah S, Sindher SB, Mallajosyula V, Gao F, Sigal N, Kowli S, Gupta S, Pellegrini K, Tharp G, Maysel-Auslender S, Hamilton S, Aoued H, Hrusovsky K, Roskey M, Bosinger SE, Maecker HT, Boyd SD, Davis MM, Utz PJ, Suthar MS, Khatri P, Nadeau KC, Pulendran B. 2021. Systems vaccinology of the BNT162b2 mRNA vaccine in humans. Nature 596:410–416.

29. Oberhardt V, Luxenburger H, Kemming J, Schulien I, Ciminski K, Giese S, Csernalabics B, Lang-Meli J, Janowska I, Staniek J, Wild K, Basho K, Marinescu MS, Fuchs J, Topfstedt F, Janda A, Sogukpinar O, Hilger H, Stete K, Emmerich F, Bengsch B, Waller CF, Rieg S, Sagar, Boettler T, Zoldan K, Kochs G, Schwemmle M, Rizzi M, Thimme R, Neumann-Haefelin N, Hofmann M. 2021. Rapid and stable mobilization of CD8+ T cells by SARS-CoV-2 mRNA vaccine. Nature 597:268–273.

30. 30. Saini SK, Hersby DS, Tamhane T, Povlsen HR, Amaya Hernandez SP, Nielsen M, Gang AO, Hadrup SR. 2021. SARS-CoV-2 genome-wide T cell epitope mapping reveals immunodominance and substantial CD8^+^ T cell activation in COVID-19 patients. Sci Immunol 6:eabf7550. https://doi.org/10.1126/sciimmunol.abf7550.

31. Cui J, Li F, Shi Z-L. Origin and evolution of pathogenic Coronaviruses. 2019. Nat Rev Microbiol 17:181–192.

32. Gonzalez-Galarza FF, McCabe A, Santos EJMD, Jones J, Takeshita L, Ortega-Rivera ND, Cid-Pavon GMD, Ramsbottom K, Ghattaoraya G, Alfirevic A, Middleton D, Jones AR. 2020. Allele frequency net database (AFND) 2020 update: gold-standard data classification, open access genotype data and new query tools. Nucleic Acids Res 48:D783–D788.

33. Rammensee H, Bachmann J, Emmerich NP, Bachor OA, Stevanović S. 1999. SYFPEITHI: database for MHC ligands and peptide motifs. Immunogenetics 50:213–219.

34. 34. Vita R, Mahajan S, Overton JA, Dhanda SK, Martini S, Cantrell JR, Wheeler DK, Sette A, Peters B. 2019. The immune epitope database (IEDB): 2018 update. Nucleic Acids Res 47:D339–D343. https://doi.org/10.1093/nar/gky1006.

35. Singh H, Raghava GPS. 2003. ProPred1: Prediction of promiscuous MHC class-I binding sites. Bioinformatics 19:1009–1014.

36. Larsen MV, Lundegaard C, Lamberth K, Buus S, Lund O, Nielsen M. 2007. Large-scale validation of methods for cytotoxic T-lymphocyte epitope prediction. BMN Bioinformatics 8:424. https://doi.org/10.1186/1471-2105-8-424.

37. Alexander J, Bilsel P, del Guercio M-F, Marinkovic-Petrovic A, Southwood S, Stewart S, Ishioka G, Kotturi MF, Botten J, Sidney J, Newman M, Sette A. 2010. Identification of broad binding class I HLA supertype epitopes to provide universal coverage of influenza A virus. Hum Immunol 71:468–474.

38. Hatcher EL, Zhdanov SA, Bao Y, Blinkova O, Nawrocki EP, Ostapchuck Y, Schaffer AA, Brister JR. 2017. Virus variation resource - improved response to emergent viral outbreaks. Nucleic Acids Res 45:D482–D490.

39. Tarke A, Sidney J, Methot N, Yu ED, Zhang Y, Dan JM, Goodwin B, Rubiro P, Sutherland A, Wang E, Frazier A, Ramirez SI, Rawlings SA, Smith DM, Antunes RS, Peters B, Scheuermann RH, Weiskopf D, Crotty S, Grifoni A, Sette A. 2021. Impact of SARS-CoV-2 variants on the total CD4+ and CD8+ T cell reactivity in infected or vaccinated individuals. Cell Rep Med 2:100355. https://doi.org/10.1016/j.xcrm.2021.100355.

40. Bugawan TL, Mack SJ, Stoneking M, Saha M, Beck HP, Erlich HA. 1999. HLA class I allele distributions in six Pacific/Asian populations: evidence of selection at the HLA-A locus. Tissue Antigens 53:311–319.

41. Tokunaga K, Ishikawa Y, Ogawa A, Wang H, Mitsunaga S, Moriyama S, Lin L, Bannai M, Watanabe Y, Kashiwase K, Tanaka H, Akaza T, Tadokoro K, Juji T. 1997. Sequence-based association analysis of HLA class I and II alleles in Japanese supports conservation of common haplotypes. Immunogenetics 46:199–205.

42. Takagi A, Matsui M. 2021. Identification of HLA-A*02:01-restricted candidate epitopes derived from the nonstructural polyprotein 1a of SARS-CoV-2 that may be natural targets of CD8^+^ T cell recognition *in vivo*. J Virol 95:e01837–20.

43. Boucherma R, Kridane-Miledi H, Bouziat R, Rasmussen M, Gatard T, Langa-Vives F, Lemercier B, Lim A, Berard M, BenMohamed L, Buus S, Rooke R, Lemonnier FA. 2013. HLA-A*01:03, HLA-A*24:02, HLA-B*08:01, HLA-B*27:05, HLA-B*35:01, HLA-B*44:02, and HLA-C*07:01 monochain transgenic/H-2 class I null mice: Novel versatile preclinical models of human T cell responses. J Immunol 191:583–593.

44. Foung SKH, Taidi B, Ness D, Grumet FC. 1986. A monoclonal antibody against HLA-A11 and A24. Hum Immunol 15:316–319.

